# Observing an antisense drug complex in intact human cells by in-cell NMR

**DOI:** 10.1101/589812

**Authors:** Judith Schlagnitweit, Sarah Friebe Sandoz, Aleksander Jaworski, Ileana Guzzetti, Fabien Aussenac, Rodrigo J. Carbajo, Elisabetta Chiarparin, Andrew J. Pell, Katja Petzold

**Affiliations:** Department of Medical Biochemistry and Biophysics, Karolinska Institute, Stockholm, Sweden; Department of Materials and Environmental Chemistry, Arrhenius Laboratory, Stockholm University, Stockholm, Sweden; Bruker BioSpin, Wissembourg, France; Analytical & Structural Chemistry, Oncology, IMED Biotech Unit, AstraZeneca, Cambridge, UK

**Author notes:** these authors contributed equally, co-first authors (Judith Schlagnitweit, Sarah Friebe Sandoz).

**Keywords:** NMR, nucleic acids, in-cell NMR, DNP, antisense oligonucleotide

## Abstract

Gaining insight into the uptake of drugs into cells, trafficking and their target engagement enhances understanding of the drug’s function and efficiency. Here we study an antisense oligonucleotide drug (ASO) delivered into HEK293T and HeLa cells, by Nuclear Magnetic Resonance (NMR). Using a combination of transfection, cryoprotection and dynamic nuclear polarization (DNP), we were able to detect the drug directly in intact frozen cells. Activity of the drug was confirmed by qRT-PCR, measuring downregulation of its target mSTAT3. Applying DNP NMR to frozen cells, we overcome limitations of traditional solution-state in-cell NMR (e.g. size, stability and sensitivity) as well as of visualization techniques, where (e.g. fluorescent) tagging of the ASO decreases its activity. The possibility to study an untagged, active drug, interacting in its natural environment, will increase insights into molecular mechanisms of delivery, intracellular trafficking and target engagement in intact cells.

---

Understanding physiological processes in detail and their inhibition by drugs is an important topic of research. High-resolution structural techniques allow us to study complex biomolecular systems, however, these methods are mostly applicable to *in vitro* samples and cannot recapitulate cellular context, indicating that these structures may differ from those in living cells. In contrast, functional data is routinely obtained in tissues or cells using visualization techniques, e.g. confocal microscopy that requires tagging for visualization by chemical modification of the molecule of interest. Thus, while the context in which data is acquired is highly relevant, the tagged system may exhibit altered behaviour, e.g. cellular trafficking and/or function. Therefore, a cellular structural biology approach is needed, combining the advantage of the biologically relevant context of the cell with an atomic-resolution technique.

In this work, we studied danvatirsen, a high-affinity 16-nucleotide synthetic antisense oligonucleotide (ASO) with a gapmer design and a phosphorothioate (PS) backbone throughout the sequence^[1]^ (SI 1.1 and Figure S01). The phosphorothioate backbone is standard to increase resistance to degradation by nucleases while activation of RNaseH1 remains intact^[2]^. Danvatirsen downregulates the mRNA of the transcription factor STAT3, and is currently in clinical trials showing antitumor activity in lymphoma, non-small cell lung cancer^[1]^ and non-Hodgkin’s lymphoma^[3]^. However, understanding of intracellular trafficking and targeting processes is currently a bottleneck in ASO drug development^[4,5]^, and until now no reliable method allows the study of oligonucleotide structures in their active macromolecular complexes in human cells.

First, confocal microscopy images of transfected ASO in human cells were acquired. A biotin tagged ASO was coelectroporated (2% of 12.5 μM) with unmodified ASO into HEK293T cells and imaged by fluorescently labeled streptavidin (Figure 1a&b). The biotinylated ASO concentrates in granular structures inside the cell (transfection rates SI 1.9), likely reflecting accumulation of the drug in the endocytic pathway or in cytoplasmic structures, as supported by literature^[6]^. We further tested a fluorescent Cy3 tag of ASO for free uptake (100% of 1.25 μM), as the Cy3 label is more sensitive and produces more dispersed signals than biotin. After incubation of HeLa cells with Cy3-labeled ASO, all cells had taken up ASO. Again, ASO appears in dotlike structures, but is more dispersed in the cell than for electroporated cells (Figure 1c&d). Interestingly, these chemically tagged ASOs have significantly worse downregulation properties compared to regular non-tagged ASO, shown by qRT-PCR for pharmaceutically relevant doses of 1.25μM (67.9±5.1% for non-modified ASO, 34.9±8.9% for biotin-labeled and 37±17% for Cy3-labeled ASO, Figure 2e, details in SI 1.9, 1.10, 2.1-2.3, 2.6, 2.7). As the tagged versions are significantly less efficient, they might not reflect how untagged ASO behaves in the cell, highlighting the problem that drug modifications with e.g. fluorescent or biotin tags, can alter activity or localization, and stressing the need for a tag-free technique.

**Figure 1.**
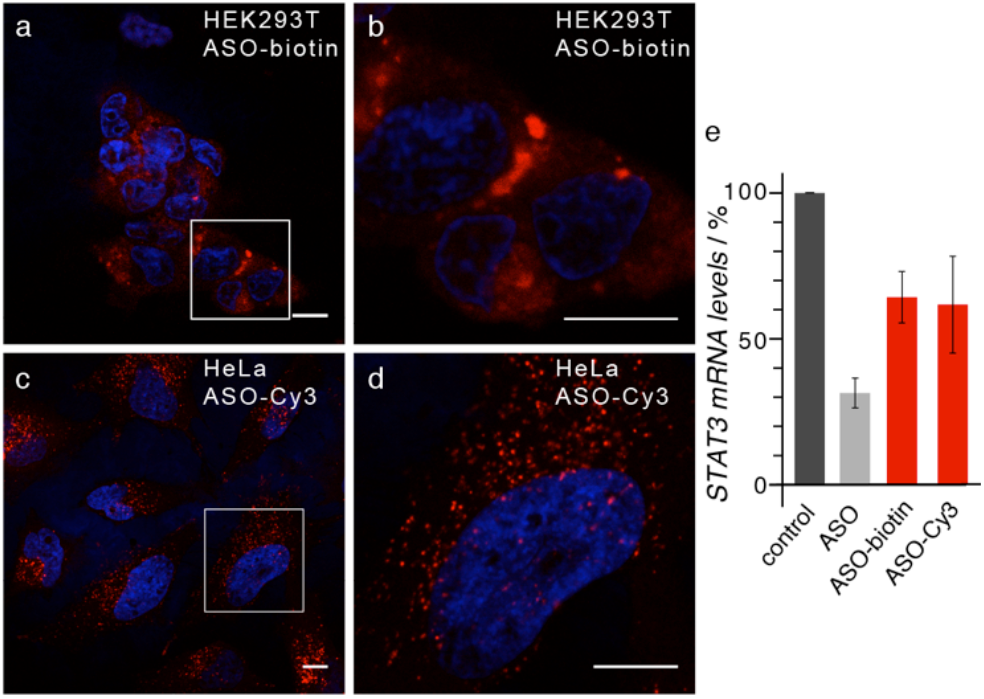
Immunofluorescence of tagged ASO correlated with activity (a) Confocal microscopy of HEK293T electroporated with 12.5 μM ASO containing 2% biotinylated ASO (blue: Hoechst 33528-nucleus, red: biotinylated ASO-AF647-streptavidin). Scale bar is 10 μm. (b) Magnification (x4.5) indicated by white box. (c) Confocal microscopy of HeLa cells after free uptake with 1.25 μM Cy3-labelled ASO (blue: Hoechst 33528-nucleus, red: ASO-Cy3). Scale bar is 10 μm. (d) Magnification (x3) indicated by white box. (e) Comparison of STAT3 mRNA downregulation efficiency of untagged ASO, biotinylated ASO and Cy3-tagged ASO by qRT-PCR after free uptake of 1.25 μM ASO solutions in HeLa cells. n=3 (except for ASO-Cy3: n=2), results show average and 1 standard deviation (s.d.). (Details see SI 1.9, 1.10 and Materials and Methods)

In-cell NMR is a tag-free technique and allows the study of structural properties at atomic resolution of, e.g. proteins^[7–13]^ and small DNA/RNA hairpins^[14–17]^ in cells. Therefore, ASO was delivered into human embryonic kidney cells (HEK293T) by electroporation, as used previously for solution-state in-cell NMR studies^[9,17]^, to ensure robust uptake, and a classical in-cell solution-state NMR spectrum was acquired. However, in the ^31^P NMR spectrum no ASO signal could be detected (Figure 2a). A reference spectrum (Figure 2h) shows peaks at ~55 ppm, characteristic chemical shifts for a phosphorothioate backbone (SI 1.1). In addition to the standard control for in-cell NMR, confocal microscopy on a tagged subset (see above), STAT3 mRNA levels were measured by qRT-PCR, showing significant downregulation of 97.12±0.87%, confirming functional delivery and productive uptake of the drug into the cell (Figure 2d) by electroporation. Based on these results we hypothesized that ASO forms large complexes leading to signal broadening and signal loss for solution-state NMR. This is in accordance with literature, where ASO is expected to be in complexes with the target mRNA and RNaseH1 (active complex) in addition to interacting with trafficking proteins or chaperones^[2,4,18]^.

**Figure 2.**
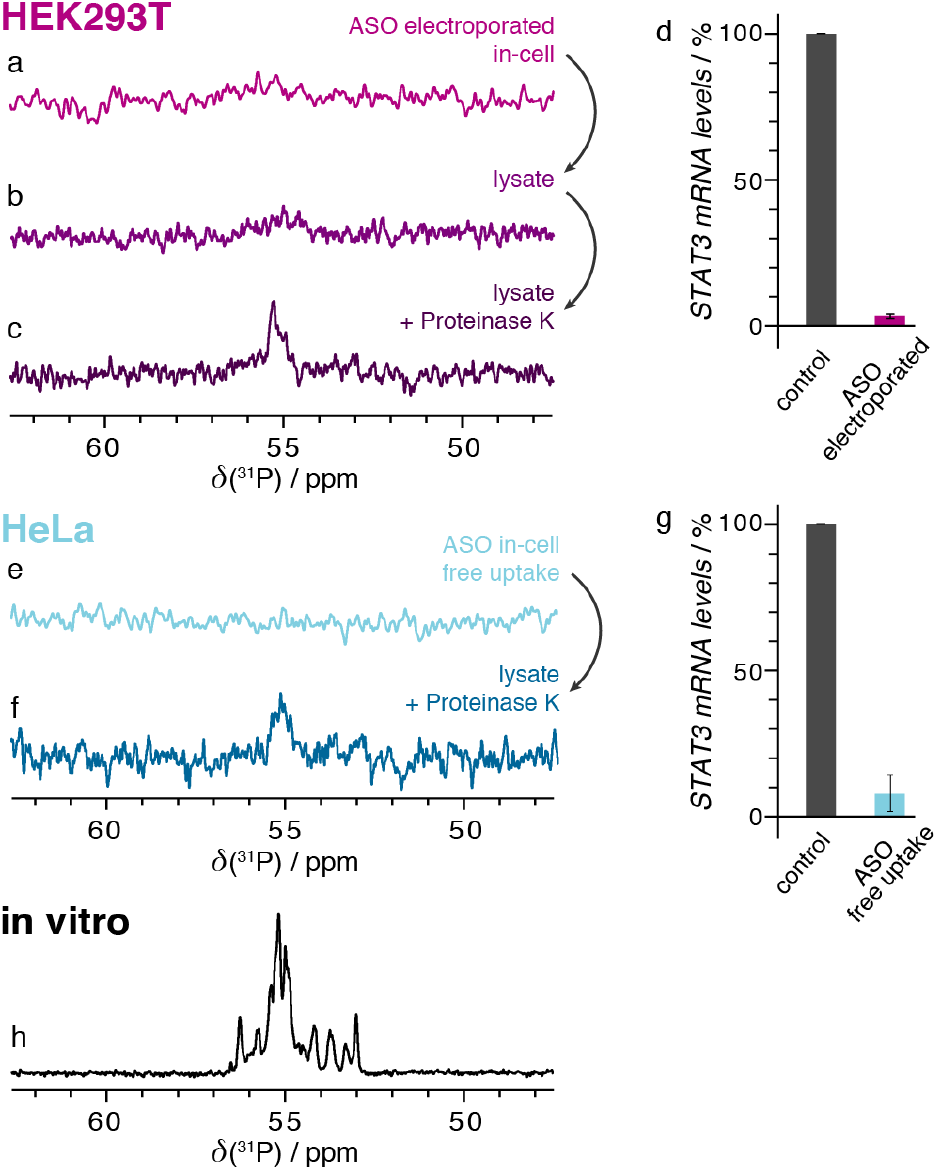
ASO solution-state NMR and activity. (a) Solution-state ^31^P in-cell NMR spectrum of HEK293T cells, electroporated with a 12.5 μM solution of ASO, after 8h (9346 scans). (b) Spectrum after lysing cells from (a) (time 16.5 h, 19456 scans). (c) Next, sample was treated with proteinase K, releasing ASO from its macromolecular complex, resulting in a detectable signal (time 16h, 18944 scans). (d) Downregulation of STAT3 mRNA levels determined by qRT-PCR in HEK293T (same as (a)), n=3, results show average and 1s.d.. (e) Solution-state ^31^P in-cell NMR spectrum of HeLa cells transfected with 12.5 μM of ASO by free uptake, again no signal could be detected after ~24 h (32768 scans). (f) Spectrum obtained after cell lysis and proteinase K treatment of sample (e) after ~48 h (77824 scans) (g) Downregulation of STAT3 mRNA levels determined by qRT-PCR in HeLa cells (same as (e)). n=3, results show average and 1s.d.. (h) Reference *in vitro* solution-state NMR spectrum of 2 mM ASO in NMR buffer. Details see SI 2.4, 2.5, 2.8.

In order to prove this hypothesis, we attempted to release the ASO from its functional complexes by lysing cells of the in-cell sample (Figure 2a) using freeze-thaw cycles and sonication, resulting in no signal in the spectrum of the lysate (Figure 2b), indicating that the complexes are still intact. Next, this lysate was treated with proteinase K in order to digest the proteins^[19]^, while endogenous levels of RNases digest the mRNA target and thus free the ASO. Only after lysing and proteinase K treatment we were able to detect a signal from free ASO (Figure 2c) at the characteristic phosphorothioate chemical shift, finally demonstrating the presence of ASO in HEK293T cells to begin with. As a control, non-transfected HEK293T cells were measured with no detectable signal at 55 ppm, even after lysis and proteinase K treatment (SI 1.2). The ^31^P ASO signal obtained after cell lysis and proteinase K treatment provides further evidence that the compound was successfully incorporated into cells and is engaged in a biologically active macromolecular complex and non-specific protein complexes. These results indicate current limitations of solution-state incell NMR, restricting it to small and/or intrinsically disordered systems not interacting with complexes in the cell^[20,21]^.

Although electroporation is an efficient transfection method, for a pharmaceutically and biologically relevant sample, we investigated free uptake of ASO, meaning incorporation into cells without the aid of a transfection reagent. HeLa cells exhibited a stronger downregulation of STAT3 mRNA levels with this uptake mechanism, compared to HEK293T cells (SI 1.10, Figure S10), potentially due to a difference in productive and non-productive uptake pathways^[4,22]^. Figure 2e shows a solution-state NMR spectrum of an in-cell sample of HeLa cells detected in 24h. Similar to electroporated HEK293T, the in-cell NMR signal evaded detection while biological activity could clearly be observed from downregulation using qRT-PCR (92.3±6.3%, Figure 2g). Similar to HEK293T, the solution-state NMR signal could only be observed after cell lysis and proteinase K treatment (Figure 1f, SI SI 1.2, 2.3). The detected levels of ASO in HeLa cells after free uptake are approximately a third (~5 μM) of the levels that were freed after electroporation from HEK293T cells (~15 μM, Figure 1c, details in SI 1.7).

It is evident from these results that in order to detect ASO drugs using in-cell NMR during trafficking and/or in their active, complexed state, three challenges have to be overcome: (i) limited measurement times due to short cell life times and metabolic changes over time, paired with (ii) low (physiological) concentrations of the molecule of interest, and (iii) the inability to detect macromolecule complexes and machines by solution-state NMR^[20,21]^. The high-resolution method of choice to overcome the size limitation (challenge (iii)) is magic-angle spinning (MAS) solid-state NMR. However, cells are destroyed by MAS frequencies >4 kHz^[23]^, leading to broadened, low intensity signals, due to only partial averaging of the anisotropic spin interactions by low MAS paired with short cell lifetimes^[24,25]^. Consequently, a solid-state NMR spectrum obtained from the ASO in the HEK293T cells at 298K did not show any signals (SI 1.3).

In order to overcome the prevailing challenges (i), lifetime of cells, and (ii), low signal intensity, we applied a low-temperature solid-state NMR approach on frozen cells as shown for bacterial cells^[10,11]^. It is standard procedure in molecular biology to preserve and store human cells at low temperatures using cryoprotecting agents (e.g. 10% DMSO) to prevent cells from rupturing due to ice crystal formation. Cells stay intact for extended amounts of times and can be recultivated later, indicating a biologically relevant environment^[26]^. Furthermore, this method has the benefit of freezing cells at any chosen time point, preventing any further metabolic changes, increasing sample stability and extending the cell lifetime, allowing for nearly infinite NMR measurement times, circumventing challenge (i). Furthermore, greater stability of frozen cells allows for higher MAS frequencies enabling us to obtain usable signals from large molecular complexes, overcoming challenge (iii). Cryoprotected, frozen, transfected cells in combination with low-temperature solid-state NMR, allowed us for the first time to access structural information in the form of ^31^P chemical shifts of oligonucleotides in their functional environment within intact human cells. Figure 3a shows a ^1^H-^31^P cross polarization (CP) in-cell spectrum at 5.5 kHz MAS obtained within 25 hours on a standard solid-state NMR setup at 240 K. Electroporated HEK293T cells were prepared as for solution-state experiments (Figure 2a and SI 2.4). The ASO signal and its sidebands (dotted line and purple asterisks, respectively) can be detected, albeit with low sensitivity compared to cell background signal (~0 ppm with sidebands indicated by triangles), controls shown in SI 1.4).

**Figure 3.**
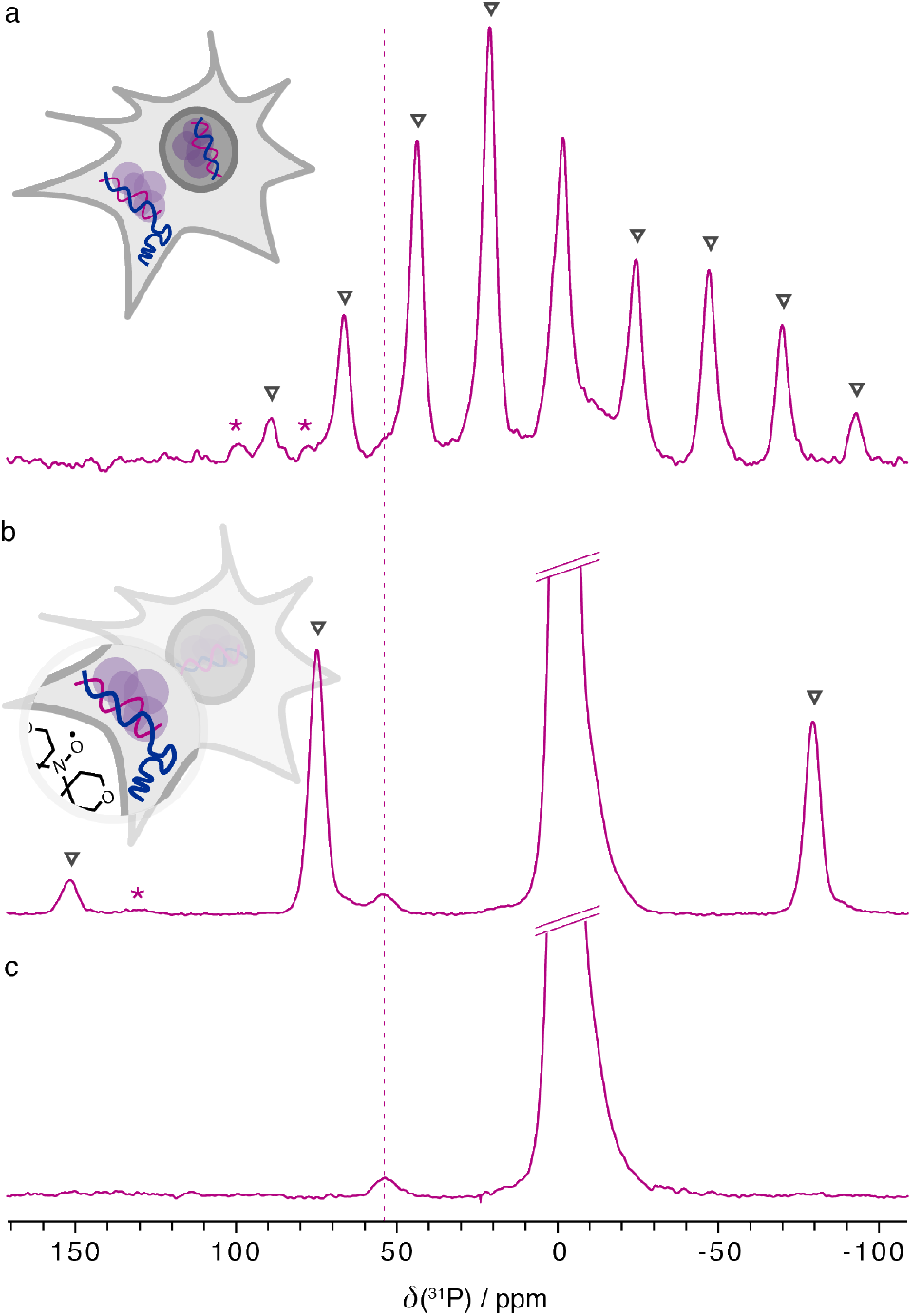
Solid-state (DNP) NMR of frozen electroporated HEK293T containing ASO. (a) ^1^H-^31^P CP in-cell spectrum obtained at 600MHz spinning at 5.5 kHz after 25h (45056 scans). The ASO signal and its sidebands are indicated by the dotted line and purple asterisks, respectively. At ~0 ppm the cell background signal is visible, leading to the sideband manifold indicated by triangles. (b) DNP enhanced ^1^H-^31^P CP in-cell spectrum of frozen cells doped using AMUPol in DMSO-d6, D_2_O and H_2_O spinning at 12.5 kHz after 1h 30min (2048 scans). The ASO signal is clearly visible at 55ppm. (c) Sum of 16 slices of a sheared DNP enhanced 2D ^1^H-^31^P CP PASS spectrum, to eliminate the sideband manifolds. Same sample as (b) but at 5 kHz MAS after 2h 40min (64 scans). Spectra in panel b and c were obtained on a 400 MHz DNP NMR spectrometer with a gyrotron frequency of 263 GHz (details in SI 2.10). This is the first time we can detect a biologically interacting oligonucleotide in intact cells.

To also truly overcome challenge (ii) of low sensitivity due to (physiological) concentrations, the frozen cryo-protected cell approach is combined with DNP leading to ICE-DNP, frozen In-Cell Enhanced NMR by DNP. DNP has been shown to improve sensitivity to detect overexpressed proteins in whole bacterial cells^[12,13]^ and obtain chemical shifts of proteins in cellular fractions^[27]^. To study the uptake of ASO we decided to follow the whole, intact cell approach and delivered the ASO either by electroporation into HEK293T, or by free uptake into HeLa cells. For DNP measurements, the freezing medium was replaced by a solution of 15 mM AMUPol in a mixture of DMSO-d6, D_2_O and H_2_O (volume ratio 6:3:1), which was added to the cell pellet leading to a ~3x dilution of the dopant solution (details in SI 2.4, 2.10). Downregulation of STAT3 mRNA for the exact samples used for DNP are shown in Figure S11. Figure 3b shows, to our knowledge, the first in-cell ^1^H-^31^P CP DNP spectrum. The spectrum was obtained from a radical-doped cell suspension of electroporated HEK293T cells at 100 K, 12.5 kHz MAS in an experimental time of only 1h 30 min, with an enhancement of 53 for the cell background signal, similar to previously reported enhancements for ^31^P of lipid samples^[28]^ (SI 1.5 for microwave-off spectra, details and controls). The increased MAS rate possible due to lower sample temperature, and the sensitivity enhancement, due to both DNP and low temperature, benefit the spectral resolution (reduced overlap of sideband manifolds) and increase sensitivity. The in-cell chemical shift (dotted line) of ASO is now clearly visible. In comparison, a DNP in-cell spectrum of free-uptake HeLa cells, where lower overall amounts of ASO in the cell are expected, ~5μM, did not result in a detectable ASO ^31^P signal after 12h of measurement even though a similar enhancement of the background signal was achieved (SI 1.6, 2.11). Here, either the sensitivity is currently too low even under DNP conditions or, untagged ASO is distributed differently within the cell upon free uptake versus electroporation and therefore the localization of the radical with respect to ASO might be suboptimal (SI 1.6). In contrast, for electroporated HEK cells, despite the complexity of the system we measured, the sensitivity enhancement is sufficiently high to permit the recording of more sophisticated experiments. In addition to the 1D CP spectrum, a 2D ^1^H-^31^P CP-PASS^[29]^ in-cell spectrum could be acquired within 2h 40min to verify the isotropic chemical shift of the bound ASO (Figure 3c).

Although solution-state in-cell NMR of transfected oligonucleotides has been carried out before^[14–17]^, the herein investigated ASO did not result in any detectable signal using this technique. This suggests that the amount of free intracellular ASO is low and instead ASO is in macromolecular complexes with mRNA and/or different proteins, as expected from literature^[2,4,18]^. To overcome the current limitations of solution-state in-cell NMR (sensitivity, stability and size) we present a frozen in-cell DNP approach. To our knowledge, this is the first time that DNP is used to detect an exogenous oligonucleotide delivered into cells, and also that a transfected antisense drug in macromolecular complexes could be detected in intact human cells. This study was carried out on the actual ASO drug compound, which was therefore untagged and unlabelled. In the absence of any ^13^C/^15^N labels we used the characteristic ^31^P chemical shift at 55 ppm of phosphorothioates. In this case the ^31^P chemical shift is not resolved or responsive enough to monitor structural changes (SI 1.1), however, this could in the future be overcome by ^13^C/^15^N labelling (which is biologically and chemically inert) on one of the bases in the ASO. Sensitivity improvements with DNP have made it possible to detect relatively low intracellular amounts of ASO after electroporation, ~15μM, though even lower concentrations of ASO after free uptake in HeLa cells evaded detection within an overnight measurement. However, with further optimization of radicals and ongoing improvement of DNP hardware, lower detection limits will be accessible in the future, approaching clinically relevant ASO doses.

The approach should be widely applicable to study previously non-detectable drugs, interacting nucleic acids, proteins and other transfected molecules in physiological, functional complexes in intact and potentially viable cells. Furthermore it is straightforward to change the investigated molecule, vary its quantity and method of delivery into a variety of different cell-lines. The frozen cells DNP approach allows time to be stopped for a functional sample, removing acquisition time limitations due to potential cell death or further metabolization as well as size limitations of the molecule of interest. Together with on-going improvements in intracellular localization of DNP radicals^[30,31]^, e.g. TOTAPOL enhanced cell wall signals^[32]^ and nuclear localization of TotaFAM^[30]^, co-localization with a transfected molecule can lead to localized enhancements of e.g. ASOs in different compartments in intact cells. We believe that this is just a preview of the potential of this method. In the future this should enable detailed insights into molecular mechanisms of a drug’s delivery and intracellular trafficking as well as its target-bound state in intact cells.

## Materials and Methods

Experimental Details see Methods in SI.

## Acknowledgements

JS acknowledges funding through a Marie Sklodowska-Curie IF (EU H2020, MSCA-IF Project No. 747446). KP acknowledges support from the Swedish Foundation for Strategic Research (Project No. FFL15-0178) and support for the 600MHz solution state NMR from Karolinska Institute. AJP acknowledges support from the Swedish Research Council (VR), grant number 2016-03441. We also thank members of the Petzold lab and the Emma R. Andersson group (Karolinska Institute) for technical support and stimulating discussions.

