## Supplementary Information for "Observing an antisense drug complex in intact human cells by in-cell NMR"

***Contents:***

**1. Additional Experimental Data**

1.1 ASO information and *in vitro* NMR data of ASO
1.2 Background/control experiments of non-transfected cells by solution-state NMR
1.3 Room temperature solid-state NMR experiments
1.4 Background/control experiments low-temperature solid-state NMR and DNP
1.5 Microwave-off experiments and enhancement factors
1.6 DNP-NMR experiments on transfected HeLa cells
1.7 Estimation of ASO concentration
1.8 Cell viability data
1.9 Microscopy data
1.10 Additional qRT-PCR data

**2. Materials and Methods**

2.1 Cell lines and culture
2.2 Transfection with electroporation
2.3 Gymnosis/free uptake of ASO
2.4 Preparing in cell NMR samples
2.5 Preparing cell lysates and proteinase K treatment for NMR measurements
2.6 RNA extraction and RT-qPCR
2.7 Biotin- and Cy3labeling and of ASO and imaging
2.8 Solution-state NMR experiments
2.9 Solid-state NMR experiments
2.10 Solid-state DNP experiments – HEK293T cells
2.11 Solid-state DNP experiments – HeLa cells

### 1. Additional Experimental Data

#### 1.1 ASO information and *in vitro* NMR data of ASO

In this work we studied danvatirsen (AZD9150), a 16-nucleotide ASO, 5'-[$^{Me}$C][$^{Me}$U][A]TTTGGATGT$^{Me}$C[A][G][$^{Me}$C]-3', with a gapmer design and a phosphorothioate (PS) backbone throughout the whole sequence. The three flanking nucleotides on either end are locked (2'-4' constrained ethyl (cET)-modified) sugars[1] (Figure S01a). These modifications stabilize the antisense oligonucleotide against degradation.[2]

Solution-state as well as low-temperature (frozen) solid-state $^{31}$P NMR spectra of a 2mM solution of ASO in NMR buffer (15 mM sodium phosphate, 0.1 mM EDTA and 25 mM NaCl at pH 6.5) were acquired for comparison and are shown in Figure S01b and c, respectively. The replacement of oxygen with sulfur leads to a significant change of the phosphorous chemical shift and transforms each phosphorous into a stereogenic center.[3] It can be seen from the solution state spectra that ASO adopts different coexisting structures corresponding to different diastereoisomers, hence for the 16 nucleotides more than 16 signals can be observed in the solution-state $^{31}$P spectrum (Figure S01b). In the frozen solid-state NMR spectra all phosphate groups contribute to the signal ~55ppm (Figure S01c).

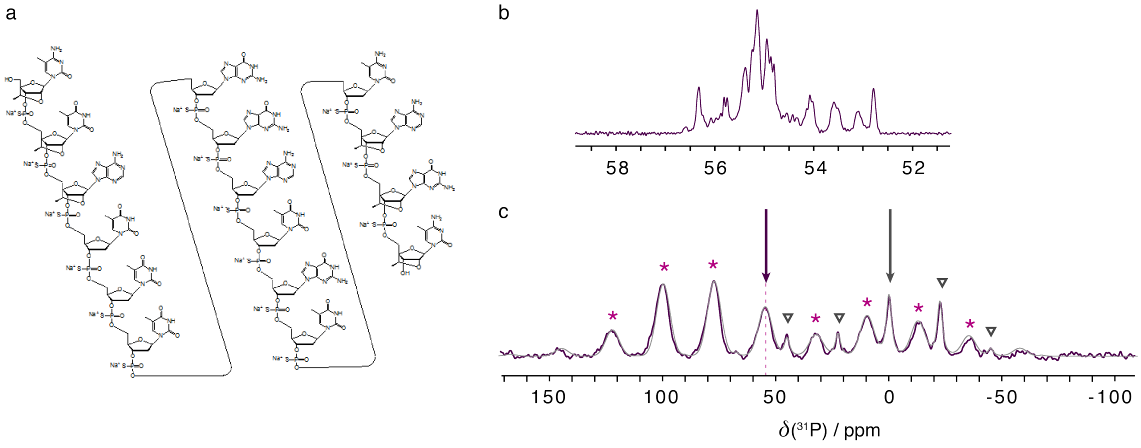


***Figure S01.*** (a) Structure including modifications of AZD9150. (b) *In vitro* $^{31}$P solution-state NMR spectrum of an ASO (danvatirsen) in buffer. 128 scans were summed up with a recycle delay of 2s (total experimental time: ~ 4min). (c) $^{1}$H-$^{31}$P CPMAS solid-state NMR spectrum obtained at 235K from a frozen ASO solution sample on a Bruker 600MHz spectrometer in a 4 mm rotor spinning at 5500 Hz. 3488 scans were summed up with a recycle delay of 3s leading to a total experimental time of ~3h. See Material and Methods section for further details. The ASO signal's centerband is visible at 55 ppm, its sidebands are indicated by purple asterisks. The narrower signal at 0 ppm, which also experiences a smaller CSA, arises from the phosphate of the NMR buffer (sidebands indicated by black triangles). The light grey line shows a Haeberlen fit obtained from Bruker TopSpin SOLA software for the two resonances indicated by black arrows (ASO: $\delta$(iso) = 56.02 ppm, $\delta$(CSA) = -113.6 ppm, $\eta$(CSA) = 0.415, and phosphate buffer: $\delta$(iso) = 1.03 ppm, $\delta$(CSA) = -60.27 ppm, $\eta$(CSA) = 0.039)

#### 1.2 Background/control experiments of non-transfected cells by solution-state NMR

Control experiments of the solution-state NMR background were carried out on non-transfected HEK293T and HeLa cells to verify that the signal at ~55 ppm solely arises from the ASO taken up by the cells. Non-transfected HEK293T cells were therefore treated exactly the same way as electroporated HEK293T cells (see Material and Methods section) and the spectrum obtained from the intact and later lysed and proteinase K treated cells, under exactly the same conditions as for the actual sample, is shown in Figure S02d. The same procedure was followed to obtain a background spectrum for non-transfected HeLa cells (Figure S02g) to compare to the ASO signal obtained after free uptake. It can clearly be seen that there is no background signal arising from non-transfected cells at ~55 ppm, hence the signal at 55 ppm can be attributed to the transfected ASO.

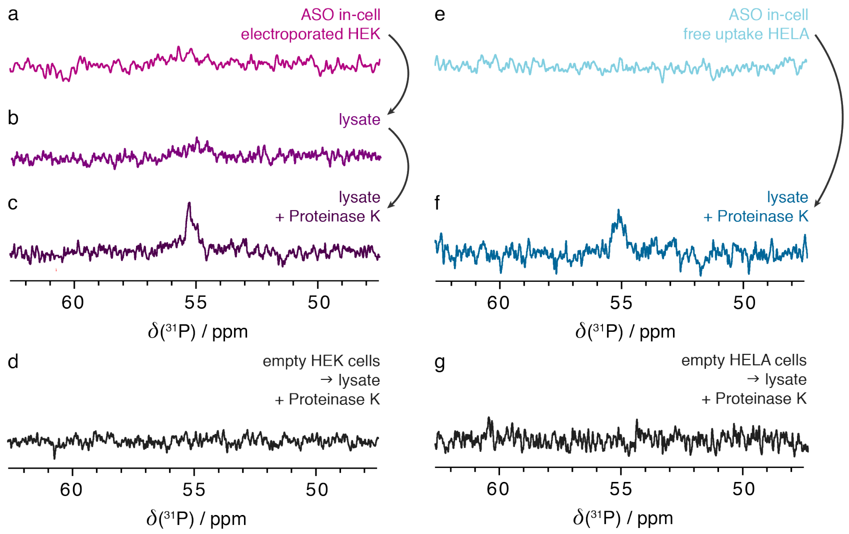


***Figure S02.*** $^{31}$P solution-state NMR spectra of electroporated HEK293T cells (panels a-c, reproduced from main manuscript Figure 2 a-c) and a control spectrum of the background of whole non-transfected HEK293T cells after cell lysis and proteinase K treatment (panel d). Similar spectra are shown for HeLa cells after free uptake (panels e, f same as main manuscript Figure 2 e, f, reproduced for comparison) and a control spectrum of the background arising from non-transfected HeLa cells after cell lysis and proteinase K treatment (panel g), obtained under exactly the same conditions as the spectrum in panel f. See Material and Methods for details on sample preparation and NMR parameters.

#### 1.3 Room temperature solid-state NMR experiments

Room temperature in-cell solid-state NMR experiments were carried out on a sample of ASO-electroporated HEK293T cells (Figure S03). The MAS rate was 3020 Hz and measurement times were short to keep the cells alive during the measurement. No ASO signal was observed (no signal at ~55ppm) after a measurement time of 4.5 hrs. This is very likely due to low concentrations combined with limited acquisition times. The observable signals are cell background signals and a signal from the phosphate buffer. See Material and Methods section for further details on sample preparation and NMR measurements.

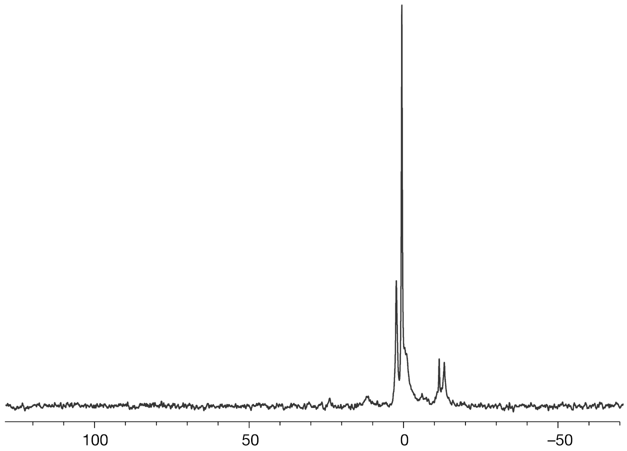


***Figure S03.*** $^{31}$P solid-state NMR spectrum obtained at room temperature from an ASO in-cell sample on a Bruker 600MHz spectrometer in a 4mm rotor spinning at 3020Hz. 8192 scans were summed up with a recycle delay of 2s, total experimental time 4.5 h. See Material and Methods section for further details.

#### 1.4 Background/control experiments of non-transfected cells by low-temperature solid-state NMR and DNP

A solid-state NMR control experiment was carried out to obtain background signals of frozen non-transfected HEK293T cells to verify that the signal at ~55 ppm solely arises from the ASO taken up by the cells. Non-transfected HEK293T cells were therefore treated exactly the same way as electroporated HEK293T cells (see Material and Methods section) and an NMR spectrum was acquired. Figure S04a shows an overlay of the spectra obtained from electroporated as well as non-transfected HEK293T cells. Especially the first and second sideband (indicated by asterisks in Figure S04a) of the ASO signal can be clearly distinguished from the background. It should be noted that this is similar to the spectrum recorded of a frozen ASO solution where the CSA also leads to a higher intensity for the sidebands than the center band (Figure S01c).
A DNP spectrum of non-transfected, radical doped HEK293T cells is compared to the spectrum obtained from electroporated cells (overlay shown in panel b). The non-transfected cell sample was prepared exactly the same way as the ASO in-cell HEK293T sample, however, the enhancement for the signal at ~0ppm of the non-transfected cells is even slightly bigger (~100). It can be seen that the background signals around -15 to -20 ppm are more intense in this sample with respect to the dominating phospholipids background signal. At 12.5 kHz MAS frequency the sidebands of those signals unfortunately overlap with the region around 55 ppm. A comparison of the spectra in the area -15 to -20 ppm shows more intense background signals for the non-transfected (grey) cells, while in the area around 55 ppm, the ASO signal of the electroporated cells has the highest intensity compared to the sidebands of the background signal. This indicates that the signal at 55 ppm in the electroporated cells arises indeed from the ASO. A second control spectrum was acquired spinning at 7.5 kHz MAS, to support the evidence that there is no background signal in non-transfected cells at 55 ppm (Figure S04c).

It can be seen in this comparison, that even though all samples were prepared using the same HEK293T cell line, the background signals in the chemical shift region around -15 to -20 ppm vary in intensity. A reason for this could be that signals in this region can arise from ATP/ADP[4] and NADH/NADPH[5] and those levels might vary for cells in different cell cycles or cells after free uptake vs cells after electroporation vs non-transfected cells.

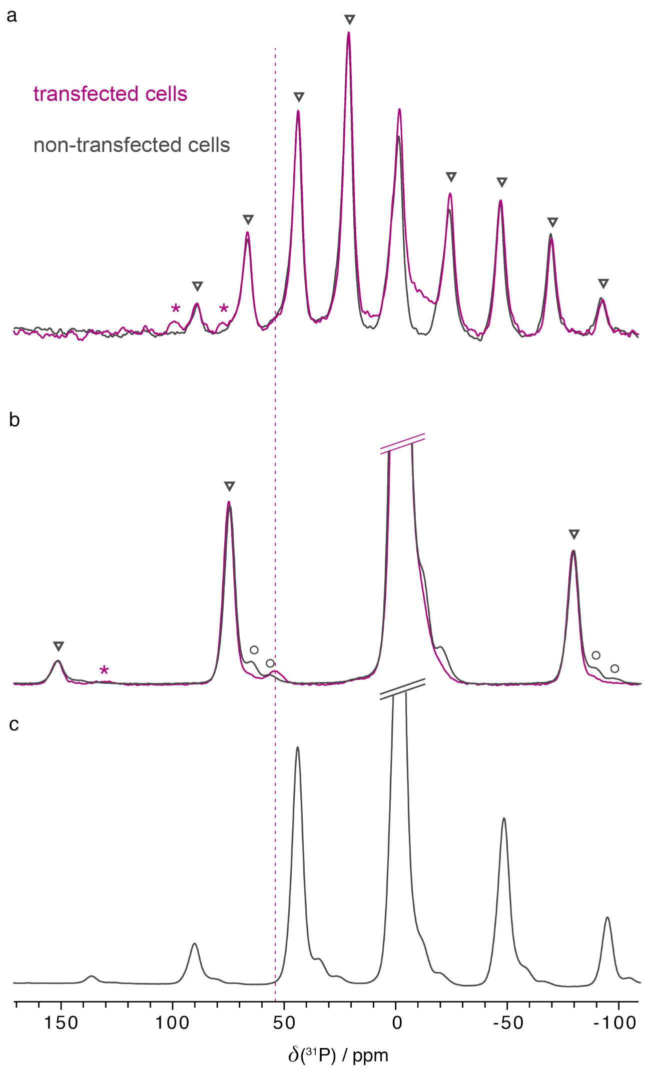


***Figure S04.*** (a) $^{1}H$-$^{31}P$ CPMAS solid-state NMR spectrum obtained at 240K from a frozen non-transfected HEK293T cell sample on a Bruker 600MHz spectrometer in a 4 mm rotor spinning at 5500 Hz. The spectrum is shown in grey. For comparison, the in-cell spectrum obtained after electroporation (same as in main manuscript Figure 3a) is also shown (purple). Sidebands of the ASO signal are indicated by asterisks, while triangles indicate sidebands of the signal arising at ~0ppm (likely phospholipids). (b) DNP enhanced $^{1}H$-$^{31}P$ CP spectrum of a frozen non-transfected cell suspension in a solution of AMUPol in DMSO-d6, $D_2O$ and $H_2O$ spinning at 12.5 kHz. The spectrum is shown in grey, sidebands of the additionally arising background signal at ~-20 ppm are indicated by grey circles. For comparison, the in-cell spectrum obtained after electroporation (same as in main manuscript Figure 3b) is also shown (purple). Sidebands of the ASO signal are indicated by asterisks, while triangles indicate sidebands of the signal arising at ~0ppm (likely phospholipids). Experimental conditions of the two overlaid spectra are exactly the same. (c) Same sample as (b) but obtained spinning at 7.5 kHz. See Material and Methods section for further details.

#### 1.5 Microwave off experiments and enhancement factors

To estimate the enhancement for the electroporated HEK293T cell sample with 15mM AMUPol in 60% DMSO-d6, 30% $D_2O$, 10% $H_2O$, $\mu W_{on}$ and $\mu W_{off}$ experiments were compared. An enhancement factor of ~53 can be determined for the cell background signal visible also in the $\mu W_{off}$ spectrum (Figure S05). This enhancement factor is similar to $^{31}P$ enhancements previously reported for lipids[6]. An enhancement factor for the ASO inside the cell cannot be determined directly. We estimate it to be slightly lower judging from the intensity of the background signal compared to the ASO signal in the enhanced spectrum and the spectrum obtained without DNP at 240K on a standard solid-state NMR set-up.

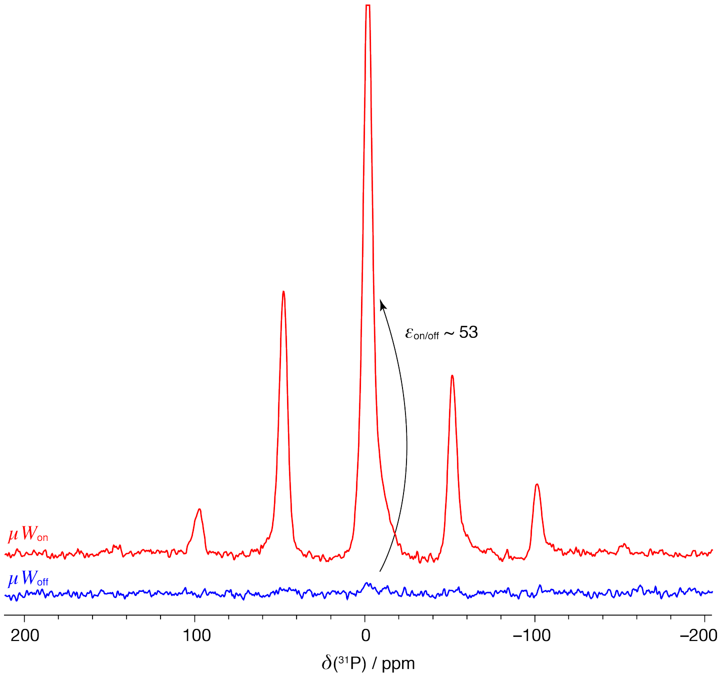


***Figure S05.*** $^{1}H$-$^{31}P$ CP solid-state NMR spectrum obtained at 97K from an ASO in-cell (electroporated HEK293T cell) sample on a Bruker 400MHz DNP spectrometer with a gyrotron frequency of 263 GHz, equipped with a 3.2mm low temperature HXY MAS probe spinning at 8000 Hz. 64 scans were summed up in both, the microwave on (red) and the microwave off (blue) spectra with a recycle delay of 2.6 s leading to a total experiment time of 3min. See Material and Methods section for further details.

#### 1.6 DNP NMR experiments on transfected HeLa cells

A free uptake HeLa in-cell sample mixed with 15mM AMUPol in 60% DMSO-d6, 30% $D_2O$, 10% $H_2O$ (preparation same as for HEK293T cells, final DMSO concentration ~20%) was prepared and $^{1}H$-$^{31}P$ CP DNP spectra were acquired. To estimate the enhancement $\mu W_{on}$ and $\mu W_{off}$ experiments were compared. An enhancement factor of ~61 can be determined for the cell background signal visible also in the $\mu W_{off}$ spectrum (Figure S06a). This enhancement factor is similar to $^{31}P$ enhancements observed in HEK293T cells.
In an attempt to detect ASO in HeLa cells, a longer CP spectrum was acquired over night (12h). 7.5 kHz MAS frequency was chosen to avoid possible overlap with the background signal with an isotropic chemical shift of -20 ppm. Despite the longer data accumulation no ASO signal could be detected of ASO in those free uptake in-cell samples.

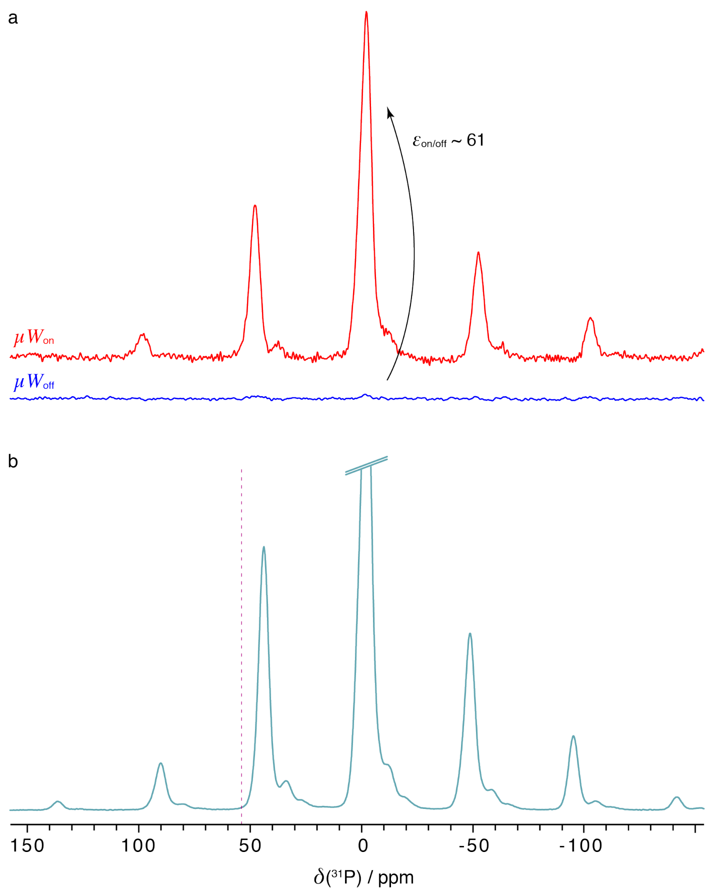


***Figure S06.*** $^{1}$H-$^{31}$P CP solid-state NMR spectra obtained at 97K from an ASO in-cell (free uptake in HeLa cells) sample on a Bruker 400MHz DNP spectrometer with a gyrotron frequency of 263 GHz, equipped with a 3.2mm low temperature HXY MAS probe. (a) MAS frequency 8000 Hz. 64 scans were summed up in both, the microwave on (red) and the microwave off (blue) spectra with a recycle delay of 2.6 s leading to a total experiment time of 3min. (b) MAS frequency 7500 Hz. 16384 scans were summed up with a recycle delay of 2.6 s leading to a total experiment time of 12h. See Material and Methods section for further details.

#### 1.7 Estimation of ASO concentration

To estimate the concentration of ASO that was taken up by the cells via electroporation or free uptake a signal-to-noise comparison with spiked lysates was carried out. Spiked lysates were chosen instead of pure buffer controls to resemble the conditions of actual samples as closely as possible. Sample one was a 2 mM ASO sample in NMR buffer mixed 1:1 with HEK293T lysate leading to the 1mM ASO "over"spiked lysate (Figure S07a). To also include the effects of proteinase K treatment, we spiked a second lysate with a known, much smaller (20µM) ASO concentration. This leads to a significantly broadened signal in the lysate (Figure S07b). Following exactly the same procedure as for the first in-cell sample (electroporated HEK293T) we treated the lysate with proteinase K to free the ASO and obtain a detectable, narrower solution state NMR signal (Figure S07c) comparable to the signal we obtained from the original in-cell samples. The signal-to-noise ratio (SINO) of those spiked spectra was compared to spectra obtained from electroporated HEK293T cells upon lysis and proteinase K treatment (Figure S07f, same as Figure 2c of the main manuscript) as well as from free-uptake HeLa cells upon lysis and proteinase K treatment (Figure S07h, same as Figure 2f of the main manuscript). Preparation of lysates as well as the procedure of proteinase K treatment is described in the Material and Methods section of the main manuscript.

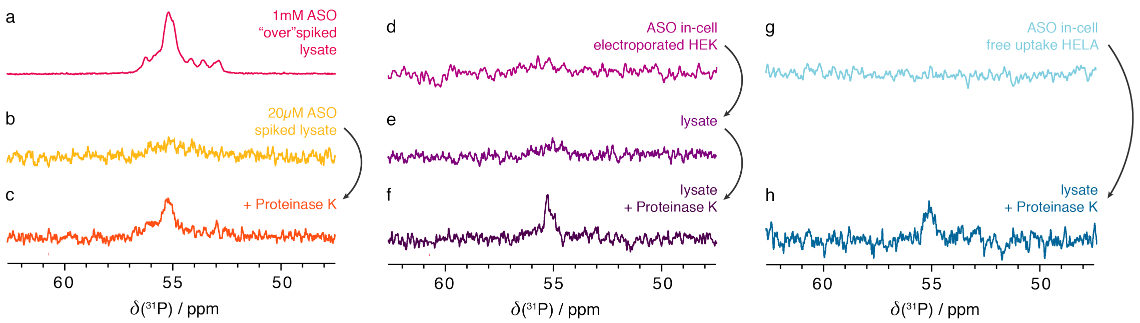


***Figure S07.*** $^{31}$P solution-state NMR spectra of spiked lysates (panels a-c) as well as of electroporated HEK293T cells (panels d-f, reproduced from main manuscript Figure 2 a-c). Similar spectra are shown for HeLa cells after free uptake (panels g,h same as main manuscript Figure 2 e,f, reproduced for comparison). See table 1 below for number of scans of the spectra of interest.

Table 1 shows ASO concentrations derived from SINO values, which were corrected for the different number of scans (NS) and then used to estimate the ASO concentration. The obtained concentrations are ~14µM for electroporated HEK293T cells and ~5µM in case of free-uptake HeLa cells. It should be noted that, given that SINO numbers are quite low, this is just a rough estimate of the amount of ASO that can be freed from the cells. This becomes apparent from the fact that the second sample was spiked to have a 20µM concentration, while the determined concentration from SINO after proteinase K treatment is lower (14µM). It should also be noted that in order to obtain a detectable signal from free-uptake cells, a more thorough proteinase K treatment (over a longer time with multiple additions of proteinase K – see Material and Methods section for details) than for electroporated HEK293T cells was necessary. The estimated uptake values in the table below are solely based on the freed ASO signal while intracellular ASO differences between HEK293T and HeLa samples may be even higher.

***Table 1:*** Calculated concentrations of freed ASO in different lysate samples based on signal to noise values obtained from solution state NMR spectra.

| **Sample** | **Spectrum** | **SINO** | **NS** | **SINO/(NS)$^{1/2}$** | **[ASO] / mM** |
|---|---|---|---|---|---|
| 1mM ASO in lysate "over"spiked | Figure S07a | 97.30 | 1024 | 3.041 | 1.070 (known) |
| 20µM ASO in lysate + proteinaseK | Figure S07c | 6.47 | 26624 | 0.040 | 0.014 |
| electroporated ASO, lysed cells + proteinase K | Figure S07f | 6.02 | 18944 | 0.044 | 0.015 |
| free uptake ASO, lysed cells + proteinase K | Figure S07h | 4.16 | 77824 | 0.015 | 0.005 |

#### 1.8 Cell viability data

Cells were tested for viability with Trypan Blue and a Countess II automated cell counter (Invitrogen) according to the manufacturer's protocol. This was done after transfection, before and/or after NMR measurements, depending on the experiment.

Viability of HEK293T cells 24h after electroporation and before NMR measurements was 68% (+/- 10%, average of three independent experiments).
Viability of HeLa cells after free uptake and before NMR measurements was assumed to be between 95% and 100%, as only stably attached cells were used for experiments.

Both HEK293T and HeLa cells were viable after solution state experiments, with varying percentages.
Around 50% of HeLa cells were alive after low temperature solid-state NMR experiments with a MAS rate of 5.5 kHz at 220 K. HEK293T cells were not viable after low temperature solid-state NMR experiments. It should be noted that experimental conditions were optimized for signal intensity and not for cell viability at the moment.
Cells that were frozen in DNP solution (see Material and Methods section), could not be revived after measurements.

#### 1.9 Microscopy data and transfection efficiency estimates

Additional microscopy images for HEK293T cells electroporated with a mixture of 98% ASO and 2% ASO-biotin (overall 12.5 μM). The non-transfected cells show background staining with AlexaFluor® 647 Streptavidin, as cells have endogenously biotinylated proteins (Figure S08a). Transfected cells however show the appearance of aggregated structures, which are absent in the negative control and clearly indicate transfected cells (Figure S08b and S08c). From these images by simply counting the total number of cells as well as the number of transfected ones, we can estimate a transfection rate of 70% (transfected cells/total number of cells counted). This agrees with data obtained on optimization experiments with a fluorescently labeled small RNA molecule, where transfection rates were assessed by flow cytometry and were at 77% (+/- 7%, average of three independent experiments).

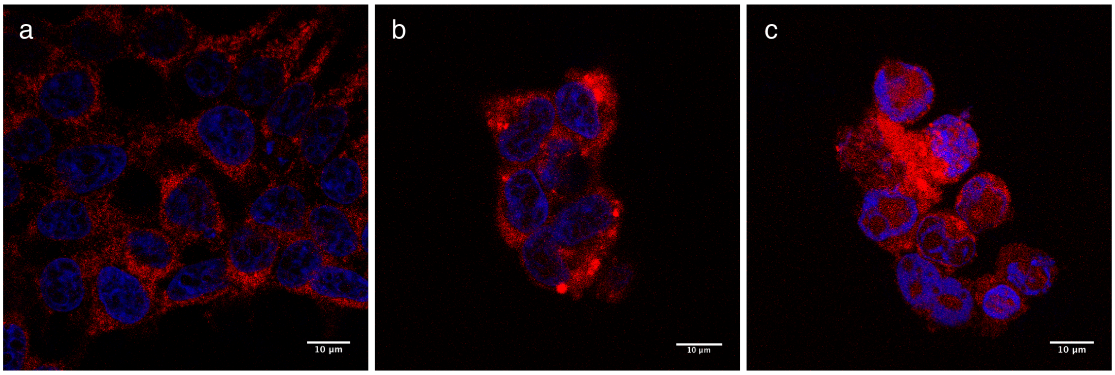


***Figure S08.*** Additional microscopy images for HEK293T cells electroporated with ASO and ASO-biotin. (a) negative control without ASO-biotin. (b) and (c) after electroporation of ASO and ASO-biotin.

Additional microscopy images for HeLa cells with free uptake of ASO-Cy3. Here, non-treated cells show no fluorescence. All cells that were imaged in ASO-Cy3 treated cells contained the drug, showing that all cells in our sample have taken up the drug via free uptake, also called gymnosis.

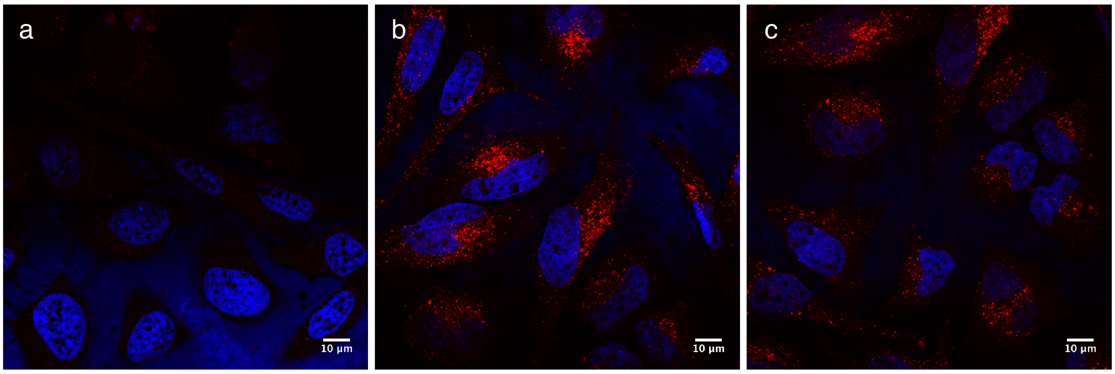


***Figure S09.*** Additional microscopy images for HeLa cells. (a) negative control without ASO-Cy3. (b) and (c) after free uptake of ASO-Cy3, different regions.

#### 1.10 Additional qRT-PCR data

Levels of STAT3 mRNA, the target of this specific ASO, were assessed by qRT-PCR and compared across cell lines and uptake mechanisms and are shown in Figure S10.

For assessment of tagged ASO activity, the ASO was labeled with two kits (see Material and Methods section for details) attaching biotin or Cy-3, respectively. The labeling might not be 100% complete and residual activity could stem from unlabeled ASO.

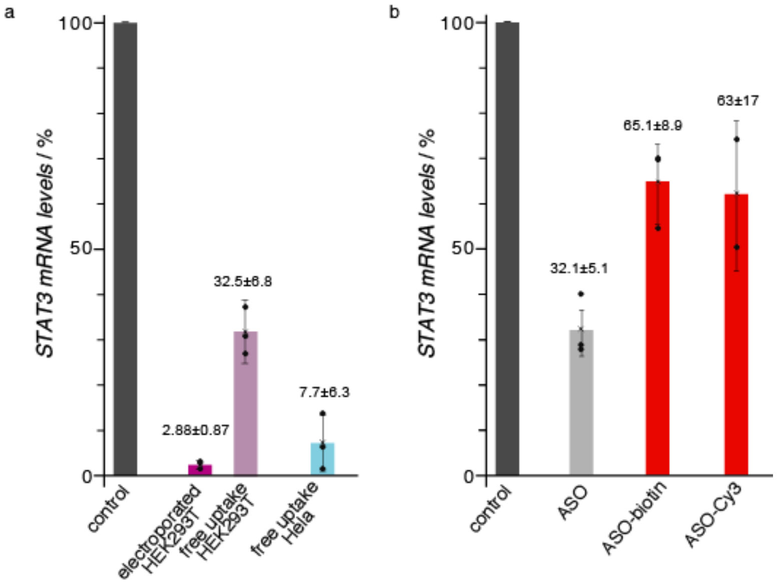


***Figure S10.*** (a) Down-regulation of STAT3 mRNA levels determined by qRT-PCR in HEK293T cells after electroporation with 1.2 mM ASO and free uptake with 12.5 µM ASO (purple bars), as well as in HeLa cells after free uptake with 12.5 µM ASO. These concentrations represent transfection concentrations as used for samples for in-cell NMR experiments. This data was also reproduced in the main manuscript Figures 1d and 1g. (b) Comparison of down-regulation of STAT3 mRNA levels determined by qRT-PCR in HeLa cells upon free uptake with 1.25µM (10-fold reduced) ASO, ASO-biotin and ASO-Cy3. These are similar concentrations as used in free-uptake studies carried out on AZD9150.[1] This data was also reproduced in the main manuscript Figure 1e.

Samples that were used for the DNP measurements were individually measured and mRNA levels of these specific experiments (electroporated for HEK293T cells, for DNP spectrum see main manuscript Figure 3, and free uptake for HeLa cells, DNP spectrum in Figure S06) are shown below. While triplicate repeats are not feasible at this point, these values compare well with the triplicate values displayed in Figure S10a.

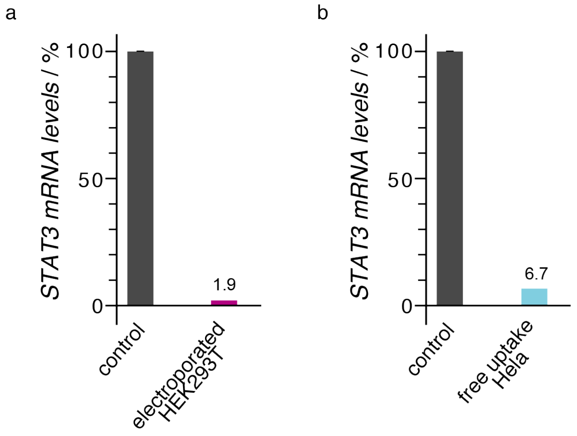


***Figure S11.*** Down-regulation of STAT3 mRNA levels determined by qRT-PCR in HEK293T cells after electroporation with 12.5µM ASO (a) and in Hela cells after free uptake with 12.5µM ASO (b) on DNP samples.

### 2. Materials and Methods

#### 2.1 Cell lines and culture

HEK293T (CRL-11268) and HeLa (CCL-2) cells were obtained from ATCC and cultured in Dulbecco's modified essential medium (DMEM, Gibco) supplemented with 10% fetal calf serum (FCS, Gibco) at 37°C and 5% $CO_2$ in a humidified incubator. Cells were used for experiments between passages 2 and 15.

#### 2.2 Transfection using electroporation

For electroporation, HEK293T cells were grown to 70-90% confluency. Per 2mm cuvette, 5*106 cells and 0.8 mg ASO were used. Cells were resuspended in 100µl Ingenio Electroporation Solution (Mirus) (leading to 1.25 mM ASO concentration in the electroporation buffer) and transfected with program A-023 in an AMAXA II electroporator (Lonza). One single pulse was applied per cuvette. Cells were resuspended immediately in warm medium (corresponding to a final 12.5 µM ASO concentration) and grown for another 24h at 37°C and 5% CO2 before being used for NMR measurements or RNA extraction.

#### 2.3 Free uptake of ASO

HeLa and HEK cells were grown to 70-90% confluency. Then, the medium was changed to medium with 12.5mM of ASO and grown for another 24h at 37°C and 5% $CO_2$ before being used for NMR measurements or RNA extraction.

It should be noted that the ASO concentrations for both transfection methods (electroporation as well as free uptake) were chosen so that it would lead to the same ASO transfection concentration with respect to a given number of cells.

#### 2.4 Preparing in cell NMR samples

Cells were washed twice with PBS and detached with trypsin. After pelleting the cells by centrifugation for 3 min at 300g and room temperature (RT), the pellet was washed with Leibovitz's L-15 medium (Gibco), supplemented with 10% FCS, resuspended in 250ml and measured in a 5mm Shigemi tube for solution state NMR. The cells were centrifuged to the bottom of the tube using a hand-cranked centrifuge. For solid-state NMR experiments at room temperature, the cells were treated as above but the cell pellet was transferred to a 4mm rotor. For solid-state low-temperature NMR experiment on frozen cells, cells were treated as above, but then frozen in 80ml freezing medium with 10% DMSO, 70% DMEM and 20% FCS in a 4mm rotor. For DNP measurements, cells were treated as above, washed twice with PBS at RT and then once with 100 ml of DNP solution (15 mM AMUPol in 60% DMSO-d6, 30% $D_2O$, 10% $H_2O$) at 4°C before resuspending in 25 ml DNP solution and loading in a 3.2mm sapphire rotor capped with a silicon insert. It should be noted that the volume ratio between cell pellet and DNP solution was ~2/3. The final DMSO concentration in the sample was therefore reduced to ~20%. Samples were frozen at -80°C for 16h before transfer to liquid nitrogen.

#### 2.5 Preparing cell lysates and proteinase K treatment for NMR measurements

For lysate from an in-cell sample, the sample was taken from the Shigemi tube and washed twice with NMR buffer and resuspended in 100 ml of NMR buffer. Cells were lysed by 5 cycles of freeze-thaw in an ethanol/dry ice bath and 3 rounds of 20s sonication in a water bath. After lysis, cells were centrifuged at 10.000g at 4°C for 10 min and the supernatant was kept for further measurements.

For proteinase K treatment, 5 ml of recombinant protein (20mg/ml, Roche) were directly added to the sample and incubated for 1h at 37°C and 15 min at 55°C.

Free uptake samples were additionally incubated for 5 days at RT and treated with proteinase K a second time before measuring. In this case both, the supernatant as well as the cell debris were kept after lysis and treated with proteinase K.

#### 2.6 RNA extraction and RT-qPCR

mRNA extraction was done with the Monarch Total RNA Miniprep Kit (NEB) according to the manufacturer's protocol. RNA concentrations were determined with a Nanodrop and 500 ng were reverse transcribed with the Maxima First Strand cDNA Synthesis Kit (Thermo Scientific).

qRT-PCR was performed with Fast SYBR™ Green Master Mix (Thermo Scientific) on a 7500 Fast Real-Time PCR System (Applied Biosystems).

Primers were the following:

| **Target** | **Forward primer** | **Reverse primer** |
|---|---|---|
| STAT3 | CGGAAGAGAGTGCAGGATCT | CAGCTCACTCACGATGCTTC |
| GAPDH | GCTCTCTGCTCCTCCTGTTC | ACGACCAAATCCGTTGACTC |
| Tubulin | TACCTTGAGGCGAGCAAAAA | TCACTGATCACCTCCCAGAA |
| Actin | CCAACCGCGAGAAGATGA | TCCATCACGATGCCAGTG |

Results were analyzed with the $DDC_T$ method. Results show the average and standard deviation of three independent biological replicas, which were run with three technical replicas each. Samples were normalized to non-treated cells, which were set to 100%.

#### 2.7 Biotin-labeling and Cy3-labeling of ASO and imaging

Biotin-labeling of the ASO was done with the Photoprobe® Biotin for Nucleid Acid Labeling Kit (Vector Laboratories), according to the manufacturer's protocol. Cy3-labeling of the ASO was done with the LabelIT® siRNA Tracker Intracellular Localization Kit (Mirus), according to the manufacturer's protocol. Cells were then treated as described above for free uptake and electroporation before RNA extraction or immunofluorescence (IF) staining.
For immunofluorescence (IF) microscopy, cells were washed twice with PBS and then fixed for 20 min with 4% paraformaldehyde at RT. Cells were washed again twice with PBS and permeabilized with 0.1% Triton-X (Thermo Scientific) for 4 min at RT. Cells were washed with PBS containing 10% FCS and blocked for 16h at 4°C.

Staining was done for 1h at RT with AlexaFluor® 647 Streptavidin (BioLegends, 0.5mg/ml) at 1:500 and Hoechst 33258 (Thermo Scientific, 20mM) at 1:1000 in PBS-FCS. Cells were washed twice with PBS and once with $H_2O$ before mounting in Mowiol mounting medium.

Images were acquired on a LSM880 (Zeiss) using Zen software. Images were processed with Fiji.

#### 2.8 Solution-state NMR experiments

Experiments were carried out on a Bruker Avance III spectrometer operating at 600 MHz ($^{1}$H resonance frequency) and equipped with a 5mm QCI ($^{1}$H/$^{13}$C/$^{15}$N/$^{31}$P) cryoprobe. NMR spectra were acquired and processed using Bruker TopSpin 3.2 and 3.5 software, respectively.
All solution state experiments shown in the manuscript are $^{31}$P spectra with direct excitation (6.9 kHz excitation) and proton decoupling (1.8 kHz). The recycle delay was set to 2s for all experiments and the carrier frequency was set to 60 ppm.

For the *in vitro* sample, a 2mM solution of ASO in NMR buffer (15 mM sodium phosphate, 0.1 mM EDTA and 25 mM NaCl at pH 6.5), 128 scans were recorded (Figure S01a).

For the HEK293T in-cell sample 9346 scans were summed up, total experimental time 8h. The data was recorded over night, in smaller chunks, interleaved with $^{1}$H spectra to monitor any changes due to variations in the sample before summing up all recorded data sets. In addition to the $^{31}$P spectra obtained with a carrier frequency set to 60 ppm, separate spectra with different carrier frequencies scanning -20 to 80 ppm were carried out to obtain information on possible signals over a broader range of frequencies. This is due to the limited excitation bandwidth of the $^{31}$P hard pulse available on our cryoprobe. No signals were observed besides cell background signals around 0 ppm, which resembled the ones observed in the slow-spinning solid-state NMR spectrum shown in Figure S03. For the HEK293T lysate before and after proteinase K treatment sample degradation was not an issue, so for the lysate 19456 scans (16.5h) and for the proteinase K treated lysates (both, electroporated and non-transfected) 18944 scans (16h) were added up. For the HeLa in-cell experiments after free uptake 32768 scans were recorded over 24 hours, while 77824 scans (48h) were acquired of the proteinase K treated lysate (of HeLa free uptake and control HeLa cells) to achieve a sufficiently high signal to noise ratio despite the low ASO concentration. It should be noted that compared to lysates, shorter measurement times were chosen for in-cell samples (e.g. 24h for HeLa free uptake) to ensure measurements were carried out on a majority of living cells. It can still be assumed that no in-cell signal could have been detected in the double amount of time (e.g. 48h) judging from the low signal to noise level of the signal after proteinase K treatment.

#### 2.9 Solid-state NMR experiments

Experiments were carried out on a Bruker Avance III spectrometer operating at 600 MHz ($^{1}$H resonance frequency) and equipped with a 4mm HXY probe. NMR spectra were acquired and processed using Bruker Topspin 3.2 and 3.5 software, respectively.

Slow MAS room temperature HR-MAS type experiments (see Figure S03) are $^{31}$P spectra with direct excitation (83 kHz excitation bandwidth). The recycle delay was set to 2s for all experiments. To guarantee cell integrity MAS frequencies were kept low (3020 Hz) and measurements were kept short. 8192 scans were summed up leading to a measurement time of 4.5 h.

For the low temperature measurements on frozen cells (Figure 2a of main manuscript) the VT gas of the probe was cooled to 220-240 K using a nitrogen heat exchanger. All $^{1}$H-$^{31}$P CPMAS experiments were acquired with a contact time of 1.5 ms and a constant $^{31}$P spin lock field of 30.4 kHz. The r.f. offset was centered at the ASO chemical shift. The $^{1}$H power was ramped from 32.5 to 39.7 kHz. 27.8 kHz Spinal-64 $^{1}$H heteronuclear decoupling was used during acquisition. MAS frequencies were kept low (5.5kHz) as well as the rf pulse powers to ensure that the cells stay frozen during the measurement. For HEK293T in-cell / background samples 45056 scans with a recycle delay of 2s were added leading to a total experiment time of 25 h. The frozen in vitro ASO spectrum (Figure S01c) was acquired under the same conditions 3488 scans were added up over the course of ~3h.

#### 2.10 Solid-state DNP experiments – HEK293T cells

Experiments were carried out on 9.4 T 263 GHz / 400 MHz Bruker Avance Neo solid-state DNP NMR spectrometer with a triple resonance 3.2 mm HXY probe. NMR spectra of frozen cells samples (electroporated as well as non-transfected HEK293T cells) doped with AMUPol were acquired at ~100K sample temperature. Spectra were acquired and processed using Bruker Topspin 4.0 software.
Figure 3b in the main manuscript shows a 1H-31P CPMAS experiment spinning at 12.5 kHz, acquired with a contact time of 2 ms and a constant 31P spin lock field of 68 kHz. The 1H power was ramped from 50 to 100 kHz. 100 kHz Spinal-64 1H heteronuclear decoupling was used during acquisition. 2048 scans with a recycle delay of 2.6 s were added leading to a total experiment time of 1 h 29 min.
Figure S05 in the supplementary shows 1H-31P CPMAS experiment acquired at 8 kHz MAS frequency with a contact time of 1.5 ms and a constant 31P spin lock field of 68 kHz. The 1H power was ramped from 53 to 107 kHz. 100 kHz Spinal-64 1H heteronuclear decoupling was used during acquisition. 64 scans with a recycle delay of 2.6 s were added leading to a total experiment time of 3 min for both, the experiment with and without continuous microwave.
In addition, a 1H-31P CP-PASS experiment spinning at 5 kHz was acquired with a contact time of 2 ms and a constant 31P spin lock field of 68 kHz. The 1H power was ramped from 48 to 92 kHz. 100 kHz Spinal-64 1H heteronuclear decoupling was used during acquisition. 224 scans with a recycle delay of 2.6 s were acquired for each slice in the 2D spectrum and 16 increments were recorded in the indirect dimension leading to a total experiment time of 2 h 40 min. The spectrum was sheared and a sum of all 16 slices is shown in Figure 3c.

#### 2.11 Solid-state DNP experiments – HeLa cells

Similar to in-cell experiments with HEK293T cells, DNP measurements were carried out on free uptake HeLa cells. Figure S06b shows a $^{1}$H-$^{31}$P CPMAS experiment spinning at 7.5 kHz, acquired with a contact time of 2 ms and a constant $^{31}P$ spin lock field of 68 kHz. The $^{1}H$ power was ramped from 50 to 100 kHz. 100 kHz Spinal-64 $^{1}H$ heteronuclear decoupling was used during acquisition. 16384 scans with a recycle delay of 2.6 s were added leading to a total experiment time of 12 h.

Figure S06a in the supplementary shows $^{1}H$-$^{31}P$ CPMAS experiment acquired at 8 kHz MAS frequency with a contact time of 1.5 ms and a constant $^{31}P$ spin lock field of 68 kHz. The $^{1}H$ power was ramped from 53 to 107 kHz. 100 kHz Spinal-64 $^{1}H$ heteronuclear decoupling was used during acquisition. 64 scans with a recycle delay of 2.6 s were added leading to a total experiment time of 3 min for both, the experiment with and without continuous microwave.
